# Functional lung imaging identifies peripheral ventilation changes in mice with muco-obstructive lung disease

**DOI:** 10.1101/2024.06.27.600946

**Authors:** Nicole Reyne, Ronan Smith, Patricia Cmielewski, Nina Eikelis, Kris Nilsen, Jennie Louise, Julia Duerr, Marcus A. Mall, Mark Lawrence, David Parsons, Martin Donnelley

## Abstract

β-ENaC-Tg mice serve as a relevant model of muco-obstructive lung disease, with impaired mucociliary clearance, mucus obstruction, chronic airway inflammation, structural lung damage, and altered lung function. The aim of this study was to undertake a comprehensive lung function and mechanics analysis of the adult β-ENaC-Tg model. β-ENaC-Tg and wild-type littermates underwent X-ray Velocimetry (XV) scans using a Permetium XV scanner (4DMedical, Melbourne, Australia). For comparative lung mechanics, lung function assessments were conducted with a flexiVent system. XV imaging demonstrated elevated ventilation defect percentage, mean specific ventilation, and ventilation heterogeneity in β-ENaC-Tg mice. Spatial analysis of ventilation maps indicated increased ventilation variability in the peripheral lung regions, as well as an increased proportion of under-ventilated areas. The flexiVent analysis indicated that compared to wild-types, β-ENaC-Tg mice have a significantly more compliant lungs with increased inspiratory capacity, reduced tissue elastance and increased hysteresivity (heterogeneity), suggesting loss of parenchymal integrity. This research highlights the utility of XV imaging in evaluating ventilation defects in the β-ENaC-Tg model and provides a comprehensive lung function analysis.

## Introduction

A group of muco-obstructive lung disorders, including cystic fibrosis (CF), chronic obstructive pulmonary disease (COPD), primary ciliary dyskinesia and non-cystic fibrosis bronchiectasis are characterised by reduced mucociliary clearance and increased mucus secretion, which leads to airway mucus obstruction (Zhou-Suckow et al., 2017). In the context of CF, a disease characterised by pathogenic variants of the cystic fibrosis transmembrane conductance regulator (CFTR) gene, mouse models fail to exhibit CF-like lung pathology (McCarron et al., 2018). To address this gap, researchers developed a mouse model overexpressing the β-subunit of the epithelial sodium channel (ENaC) under the wild-type Clara cell-specific promoter, known as β-ENaC-Tg mice (Mall et al., 2004). These mice exhibit increased sodium absorption, producing airway surface dehydration. The dehydration subsequently reduces mucociliary clearance, leads to mucus blockages, the transformation of goblet cells, and persistent inflammation in the airways (Mall et al., 2008). These mice serve as a model of muco-obstructive lung diseases.

β-ENaC-Tg mice develop early onset and chronic airway inflammation, characterised by airway neutrophil and macrophage activation (Zhou-Suckow et al., 2017, Gehrig et al., 2014, Hey et al., 2021, Trojanek et al., 2014). They develop epithelial remodelling and mucus hypersecretion, evidenced by goblet cell metaplasia and elevated airway mucins in the first 5 weeks of age. Chronic airway inflammation is associated with emphysema-like structural-lung disease, including distal airspace enlargement and alveolar wall destruction (Gehrig et al., 2014, Mall et al., 2008). Lung function studies show airflow obstruction and increased compliance, with computed tomography (CT) images showing reduced density of lung parenchyma (Zhou-Suckow et al., 2017). β-ENaC-Tg mice have also been demonstrated to develop emphysema, exhibiting increased lung volumes, distal airspace enlargement, and increased lung compliance (Mall et al., 2008).

Preclinical lung function assessments using animal models play a crucial role in advancing our understanding of the underlying mechanisms of pulmonary disease. One commonly utilised tool for assessing lung function in animal models, specifically rodents and small animals, is the flexiVent system developed by SCIREQ. This system allows researchers to conduct various lung function assessments by controlling the animal’s ventilation and applying different test perturbations (Ahookhosh et al., 2023). Functional assessments, such as those made by the forced-oscillation technique (FOT), can be delivered as either single frequency or broadband perturbations, and data fitted to the single compartment or constant phase models, respectively. The single compartment model provides assessments of total respiratory resistance and compliance, whilst the constant phase model offers a deeper understanding of airway and tissue mechanics, quantifying tissue damping, tissue elastance, and Newtonian resistance (central airway resistance) (Glaab et al., 2007). Moreover, the construction of pressure-volume loops can be used to characterise lung distensibility across the inspiratory capacity, and the use of the negative-pressure forced-expiration extension allows measurement of forced vital capacity (FVC), forced expired volume (FEV) and forced expired flow (FEF) data. This flexiVent assessment provides information about the dynamic and static properties of the respiratory system in animal models, aiding in the identification of deviations from normal lung function.

While functional assessments can provide information about the mechanical properties of the respiratory system, imaging techniques, such as CT and magnetic resonance imaging (MRI), offer researchers the ability to both visualise and quantify structural changes (Ohno et al., 2022, Wielpütz et al., 2016). These imaging modalities provide visual insights into the lung architecture and pinpoint abnormalities such as mucus plugging, air trapping, bronchiectasis and emphysema. Mathematical models of lung mechanics can provide a wealth of insights into the physical properties of the respiratory system. While some mathematical models can partition the lung’s response into airway and tissue mechanics components (Hantos et al., 1992), the spatial resolution of these models remains limited. Traditional imaging, while valuable, provides a static snapshot of the lung. Many lung diseases start locally, affecting lung parenchyma and small airways in a heterogeneous manner, with functional effects often masked by lung compensatory mechanisms until significant lung structure is compromised (Ahookhosh et al., 2023). Recent advancements have led to the development of X-ray Velocimetry (XV), which allows researchers to directly observe and quantify the movement of lung tissue during a breath, providing real-time functional ventilation data (Vliegenthart et al., 2022, Parsons and Donnelley, 2020).

β-ENaC-Tg mice were the first model used to demonstrate the translation of XV technology from a synchrotron facility (Stahr et al., 2016) towards a laboratory setting (Murrie et al., 2020, Werdiger et al., 2020). In the initial studies the authors demonstrated that measurements such as regional airflow and tissue expansion *in vivo* could be captured in live mice (Murrie et al., 2020). More recently, the development of the Permetium preclinical XV scanner (4DMedical, Australia), combined with custom XV analysis software, allows the motion of the lung tissue and therefore local ventilation to be measured in a laboratory setting. This technology reveals the complete dynamics of airflow across the lung during a single breath cycle (Dubsky et al., 2012, Fouras et al., 2012). In the period since those proof-of-principle XV studies using β-ENaC-Tg mice were performed, a full lung function characterisation of this mouse model has not been completed.

The primary study aim was to conduct a comprehensive lung function analysis of adult β-ENaC-Tg mice, employing both XV assessments and flexiVents. We hypothesised that XV would quantify and visually represent the extent and location of ventilation alterations in β-ENaC-Tg mice compared to wild-type. We expected these alterations to align with the parameterized lung mechanics obtained from flexiVent assessments.

## Results

### Histology

In this study β-ENaC-Tg and normal littermate (wild-type) mice were compared. Airway luminal mucus was detected in all β-ENaC-Tg mice, primarily in the larger and middle airways, but no mucus was found in the littermate wild-types (Figure 1). The mucus obstruction in the β-ENaC-Tg mice airways was heterogeneous, and when mucus was observed it did not fully obstruct or plug the airways.

**Figure 1:**
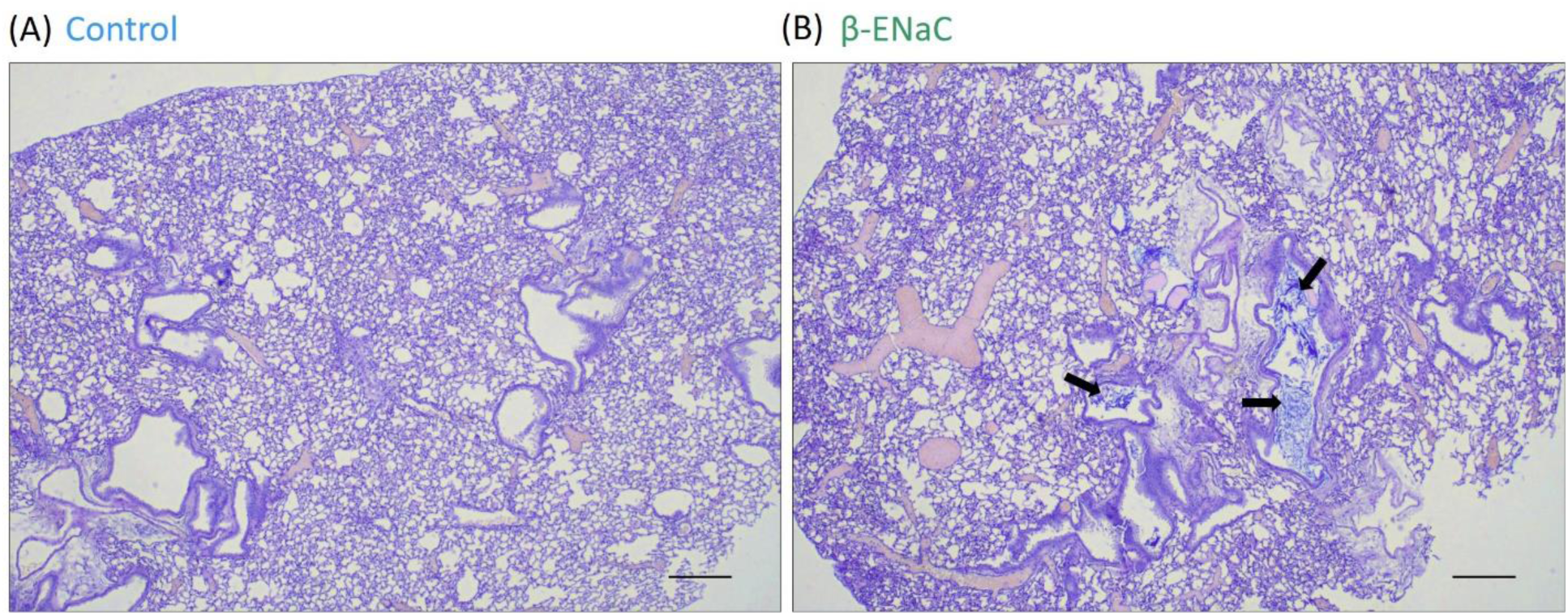
Alcian Blue/Periodic Acid Schiff staining of histological section in wild-type and β-ENaC-Tg mice. (A) wild-type mice airways were normal, compared to (B) partially mucus-obstructed airways (arrows) in β-ENaC-Tg mice. Scale bar 250 µm.

### X-ray Velocimetery (XV) imaging

XV imaging of anaesthetised and tracheostomised mice was performed using a Permetium XV scanner (4DMedical, Australia). During image acquisition mice were ventilated at a peak inspiratory pressure of 14 cmH_2_O and a positive end-expiratory pressure of 2 cmH_2_O. XV analysis showed that the normalised ventilation defect percentage, ventilation heterogeneity, and lung volume were higher in the β-ENaC-Tg mice compared to the wild-type mice, but there was no difference in mean specific ventilation (Figure 2).

**Figure 2:**
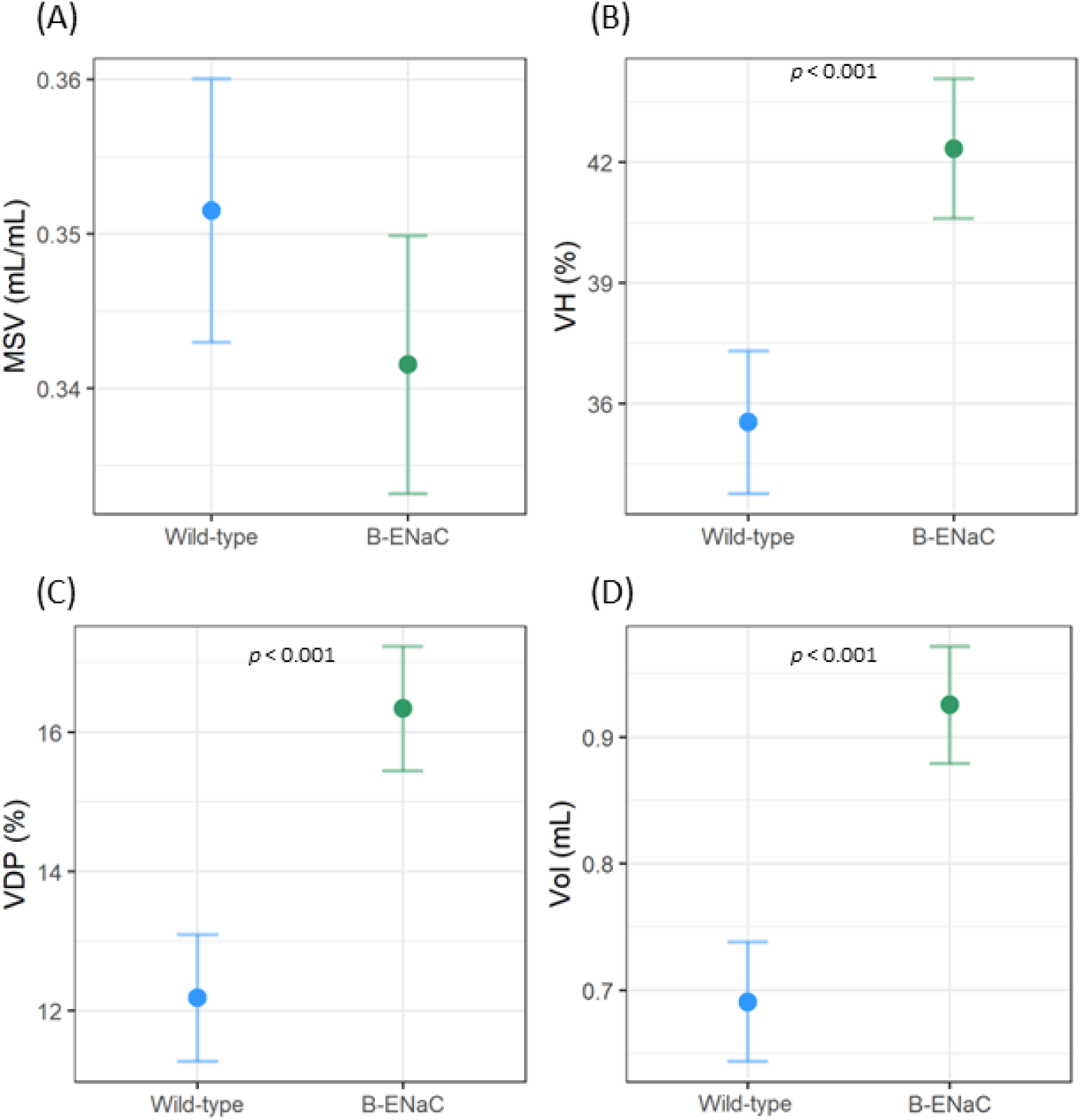
X-ray Velocimetry measurements in wild-type (blue) and β-ENaC-Tg (green) mice. (A) Mean specific ventilation (MSV), (B) Ventilation heterogeneity (VH), (C) normalised ventilation defect percentage (nVDP) and (D) the lung volume at end exhalation. (n = 13-14/genotype, standard linear regression model, estimated means and 95% CIs).

A visual examination of the XV ventilation maps indicated that the β-ENaC-Tg mice appeared to exhibit more areas of extreme high and low ventilation in the areas of the lung closest to the rib cage (Figure 3A and 3B). Based on this finding, the lungs were split into an inner core region (average 39% of total lung volume) and an outer region nearer to the rib cage. The outer and inner regions displayed obvious differences in the ventilation heterogeneity and normalised ventilation defect percentage between β-ENaC-Tg and wild-type mice (Figure 3).

**Figure 3:**
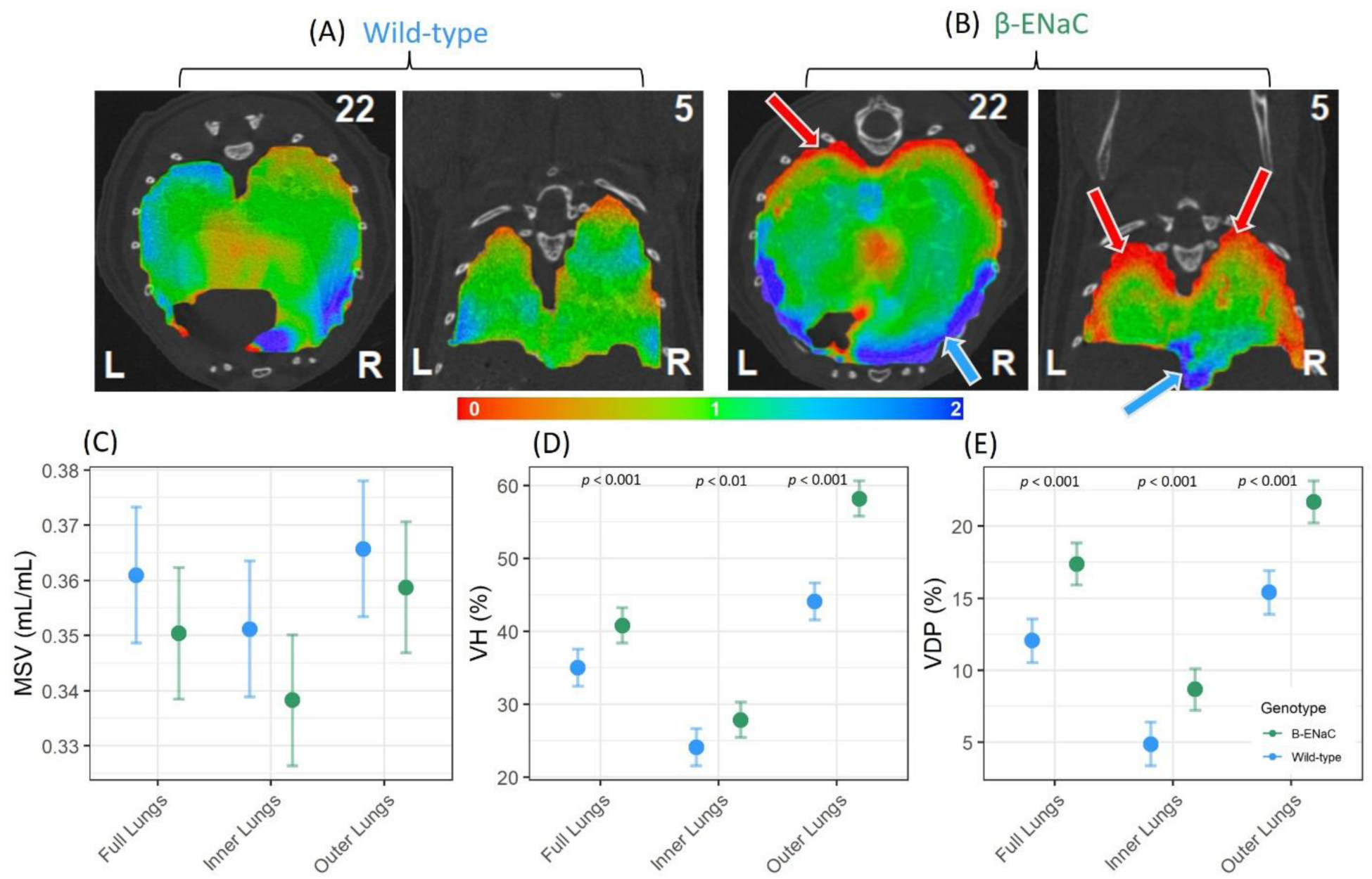
X-ray Velocimetry ventilation in the inner and outer lungs of wild-type and β-ENaC. The 2D slice mappings, derived from 3D XV volume data, from representative (A) wild-type and (B) β-ENaC-Tg mice in the transverse and coronal directions. Green indicates average ventilation, red below average ventilation and blue above average ventilation. Graphs show (C) Mean specific ventilation (MSV), (D) ventilation heterogeneity (VH), and (E) normalised ventilation defect percentage (nVDP) (n = 13-14/genotype, standard linear regression model, estimated means and 95% CIs).

### flexiVent respiratory mechanics

After XV imaging lung mechanics measurements were made using a flexiVent FX system (SCIREQ, Canada). The deep inflation manoeuvre showed that the β-ENaC-Tg mice had a higher inspiratory capacity compared to wild-types (Figure 4A), and the single frequency forced oscillation (single compartment model) demonstrated higher total respiratory system compliance (C_rs_) and lower total respiratory system resistance (R_rs_) in β-ENaC-Tg mice compared to wild-type (Figure 4B-C). The broadband forced oscillation (constant phase model) demonstrated that Newtonian (central airway) resistance (R_n_), tissue elastance (H), tissue damping (G), and tissue hysterisivity (G/H) were significantly different in β-ENaC-Tg mice compared to wild-type mice (Figure 4D-F). Average pressure volume loops constructed from mean data showed an upward shift in the pressure-volume relationship of β-ENaC-Tg mice, which was different from wild-type mice (Figure 4G-I). The static compliance (C_st_) and shape parameter K (shift in PV loop) were higher in the β-ENaC-Tg mice.

**Figure 4:**
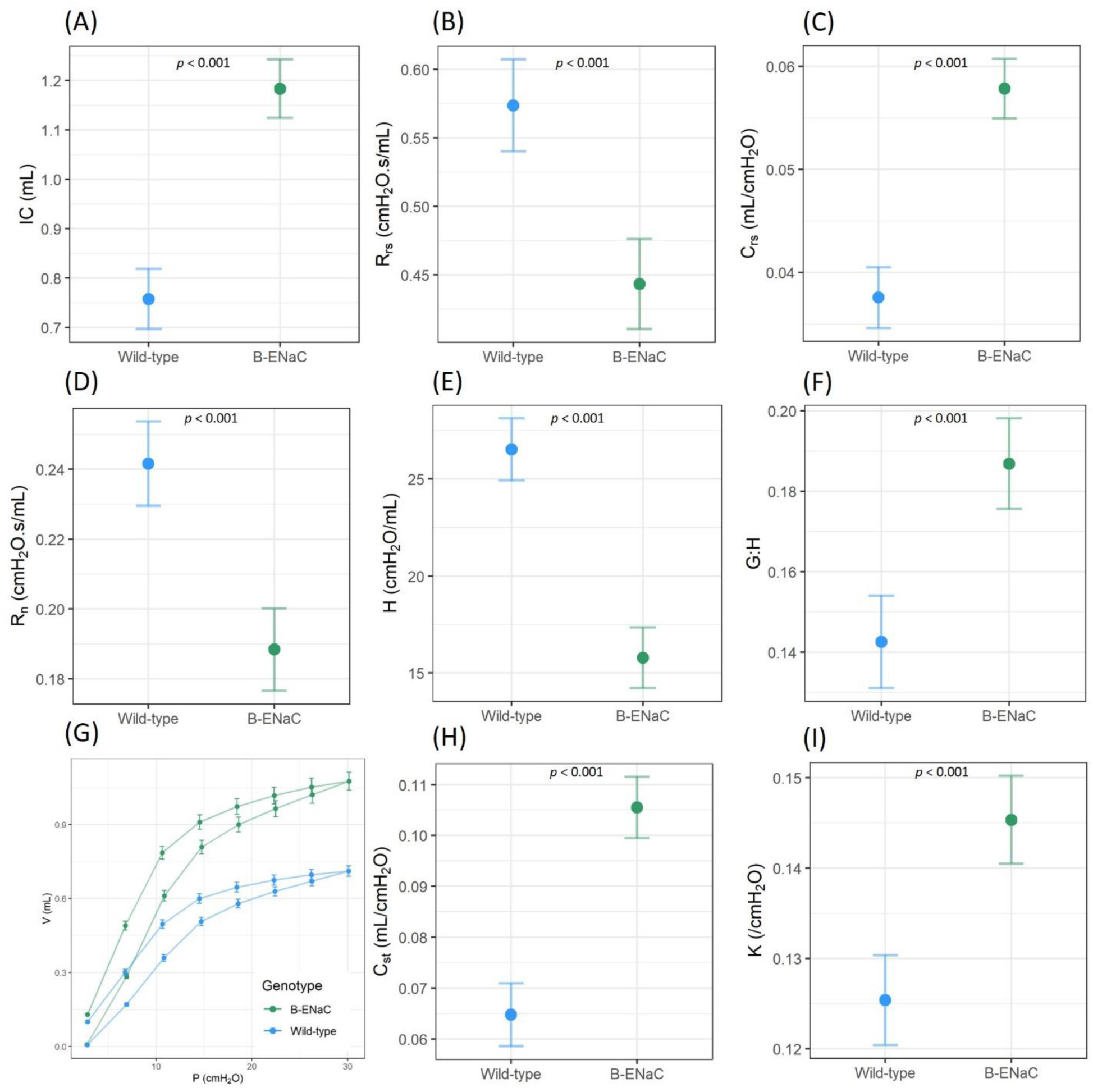
FlexiVent assessments of the respiratory system in wild-type and β-ENaC-Tg mice. Deep inflation was used to produce the (A) inspiratory capacity (IC). The single frequency forced oscillation measured the (B) total respiratory system resistance (R_rs_) and (C) total respiratory system compliance (C_rs._). The broadband forced oscillation measured the (D) Newtonian resistance (R_n_), (E) tissue elastance (H) and (F) tissue hysterisivity (G:H) of the peripheral lung compartment. The pressure volume loop (G) was used to determine the static compliance (H) and K-parameter (I) (from fitting the Salazar Knowles equation to the deflation limb of the PV loop). (n = 13-14/genotype, standard linear regression model, estimated means and 95% CIs).

When the negative pressure forced expiratory test was applied, the forced expiratory volume at 0.05s (FEV_0.05_), forced vital capacity (FVC), FEV_0.05_/FVC, and forced expiratory flow at 0.05s (FEF_0.05_) was higher in β-ENaC-Tg mice (Figure 5). The peak expiratory flow (PEF) however was not significantly different between β-ENaC-Tg mice and wild-types.

**Figure 5:**
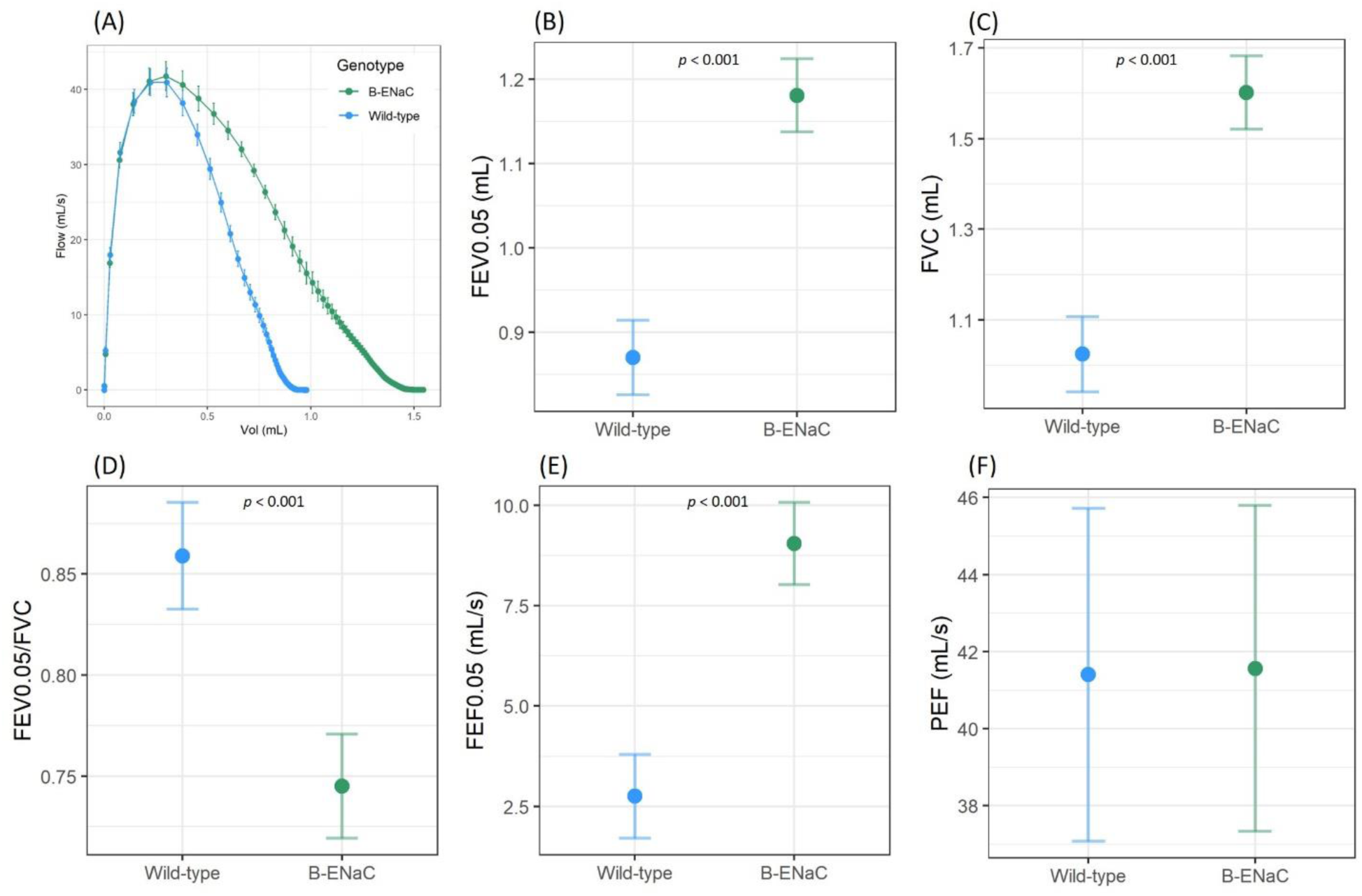
Negative pressure-driven forced expiration manoeuvres showed differences in β-ENaC-Tg mice compared to wild-types. (A) Representative mean flow-volume curves from wild-type and β-ENaC-Tg mice. (B) Forced expiratory volume (0.05 sec), (C) FVC, (D) FEV_0.05_/FVC (E) forced expiratory flow (0.05 sec), and (F) peak expiratory flow (n = 13-14/genotype, standard linear regression model, estimated means and 95% CIs).

## Discussion

Previous studies with β-ENaC-Tg mice have provided partial data on flexiVent-derived parameters (Fujikawa et al., 2020, Jia et al., 2016, Mall et al., 2008, Nakashima et al., 2022, Seys et al., 2015, Shuto et al., 2016, Stahr et al., 2016, Wagner et al., 2022, Wielpütz et al., 2011). Here, we comprehensively describe findings from a wide range of flexiVent manoeuvres for a thorough characterisation of lung function. In addition, we utilise XV to measure ventilation heterogeneity differences, and to highlight the advantages of visually representing these changes in lung function.

Forced-oscillation assessments of the β-ENaC-Tg mice – including single frequency (single compartment model) and broadband (constant phase model) methods – demonstrated mechanical properties typical of an emphysematous model: a highly compliant lung with increased lung volumes. Our results showed that β-ENaC-Tg mice have higher inspiratory capacity (from deep inflation), along with a higher total respiratory system compliance and decreased total respiratory system resistance as assessed by the single compartment model. These parameters reflect the contributions of the lung tissue, chest wall, and airways. The broadband forced oscillation (constant phase model) provided further insights by distinguishing between lung mechanics in central conducting airways and alveolar tissues. Newtonian resistance, which primarily represents the resistance of the conducting airways, was lower in the β-ENaC-Tg mice, suggesting they have larger diameter airways than the wild-type animals. Previous studies have shown that adult β-ENaC-Tg mice have enlarged distal airspaces compared to wild-type mice (Mall et al., 2008). Lower tissue damping (G) and elasticity (H) in β-ENaC-Tg mice compared to wild-types indicated a loss of structural integrity in the peripheral airways and lung parenchyma.

Additionally, a higher tissue hysteresivity (G:H) in β-ENaC-Tg mice compared to wild-types indicates lung heterogeneity, suggesting shifts in ventilation patterns across different lung regions. This finding aligns with the ventilation heterogeneity observed in the periphery of the lung in β-ENaC-Tg mice using XV imaging.

To further assess the compliance of the respiratory, pressure-driven pressure volume loops were created by inflating the lung in a stepwise manner, from PEEP (3 cmH_2_O) to 30 cmH_2_O. The β-ENaC-Tg mice displayed an upward shift in the pressure-volume relationship, higher static compliance, and K-parameter values than wild-type mice (Mall et al., 2008, Seys et al., 2015, Wagner et al., 2022). Static compliance reflects the intrinsic distensibility of the respiratory system across the entire inspiratory capacity and the K-parameter describes the curvature of the upper portion of the deflation limb of the PV loop (another representation of compliance). These results are consistent with the forced oscillation technique results, describing β-ENaC-Tg mice as having significantly more compliant lungs.

The NPFE manoeuvre generated a higher maximal inflation volume in the β-ENaC-Tg mice than in the wild-type mice, which matched the inspiratory capacity assessments. FEV0.05, FVC and FEF were higher in the β-ENaC-Tg mice, compared to the wild-type mice. Although this was consistent with previous findings (Fujikawa et al., 2020, Nakashima et al., 2022, Shuto et al., 2016), it is unusual for a high FEV to be observed in CF or emphysema patients (Wielpütz et al., 2013). Here it is likely that the increased compliance increases FVC (due to the larger inspiratory capacity) and thus FEV0.05, which is also consistent with the reduced total respiratory resistance reported by the single compartment model. Interestingly, despite histological evidence of increased mucus plugging in the β-ENaC-Tg mice, the lung function testing indicated reduced resistance and no evidence of obstruction during the NPFE manoeuvre. This is unsurprising, as it is well established that forced expiratory manoeuvres are particularly poor at measuring peripheral airway function (Cosio et al., 1978, Day et al., 2021).

A key difference between clinical spirometry data and data derived from the NPFE extension is the manner in which the data is acquired. In clinical spirometry, patients consciously and voluntarily perform exhalations, whereas NPFE assessments are made on a passive ventilated subject. Here the NPFE generates data by first inflating the lungs to 30 cmH_2_O (inspiratory capacity) and then rapidly withdrawing the air from the subject via application of -55 cmH_2_O of negative pressure to the endotracheal tube. This negative pressure application can potentially induce dynamic airway collapse and restrict airflow in a manner that cannot occur during conscious exhalation in clinical settings. Subjects with a large loss of structural tissue integrity (high compliance, i.e. emphysematous models) may be particularly prone to this effect (Shalaby et al., 2010). Despite the increased compliance, there was no evidence of any expiratory flow limitation in the β-ENaC-Tg mice, which suggests that the physiological mechanisms that result in reduced compliance are different to those typically observed in emphysema and should be taken into consideration when using this mouse as a model of emphysema.

A longitudinal study by Mall (2008) revealed that in younger β-ENaC-Tg mice, lung volumes, alveolar architecture, and alveolar size remained relatively normal, despite mucus plugging (Mall et al., 2008). However, as the mice aged, there was a subsequent increase in lung volume and distal airspace, ultimately leading to the development of emphysema. The development of emphysema in these mice is thought to be caused by several factors, including proteolytic damage of alveolar septi by proteases released from neutrophils and macrophages in chronic airway inflammation, such as neutrophil elastase (NE) and macrophage elastase (MMP12), and potentially other processes related to alveolar septation during development and the persistent hyperinflation of the lungs causing air-trapping (Mall et al., 2008). Mucus plugging, observed in previous studies with β-ENaC-Tg mice (Mall et al., 2004, Mall et al., 2008), was also observed in this study. Data here supports the association found in older people with CF, who can develop emphysema with advanced lung disease.

The normalised ventilation defect percentage was higher in β-ENaC-Tg mice than the wild-type animal, and the ventilation heterogeneity was also higher, signifying uneven or patchy airflow across the lungs caused by the CF-like muco-obstructive lung disease and emphysema phenotypes. Our findings align with the initial studies using β-ENaC-Tg mice to validate *in vivo* XV, where decreases in regional lung expansion were observed (Murrie et al., 2020, Werdiger et al., 2020, Stahr et al., 2016). Further spatial analysis of the XV ventilation maps indicated that the outer regions of the lung were more heterogeneous and exhibited a higher normalised VDP compared to the rest of the lung. This could be due to the fact that the alveolar ducts located at the edges of the lungs in β-ENaC-Tg mice are wider than those in wild-type as found in Blaskovic’s (2023) study using microCT (Blaskovic et al., 2023), or due to the mucus plugging observed in the small airways. The difference in MSV values between the inner and outer regions was more pronounced in β-ENaC-Tg mice than wild-type mice, strengthening the argument for increased heterogeneity across the whole lung. Similarly, a study on emphysema in people with CF found that CF-emphysema predominantly occurred in the subpleural regions (Wielpütz et al., 2013), the areas of the lung closest to the ribcage.

XV analysis provided spatial information to identify the origins of these lung function changes in adult β-ENaC-Tg mice. In future studies, XV could be employed to track the changing lung phenotype observed in β-ENaC-Tg mice over time, quantifying the transition from the early mucus obstructive phenotype to the later emphysematous phenotype (Mall et al., 2008). Such studies could provide valuable data for subsequent validation of XV imaging as a tool for monitoring lung disease in children with CF and other muco-obstructive lung diseases. Further, investigating the effectiveness of mucolytics on the excess mucus present in β-ENaC-Tg mice on lung ventilation changes could offer insights into how ventilation changes in the lung after successful treatment (Addante et al., 2023).

The objective of this study was to extend the characterisation of lung function and mechanics in adult β-ENaC-Tg mice and pinpoint the location and extent of airway ventilation. Here we provide a comprehensive analysis of lung function in β-ENaC-Tg mice using flexiVent and XV technology. Histology demonstrated increased mucus plugging, the flexiVent showed increased compliance and reduced respiratory resistance, and XV demonstrated increased VH and VDP. XV functional lung imaging showed that β-ENaC-Tg mice have regions of altered ventilation in the peripheral regions of the lungs.

## Methods

### Animals

All animal procedures were approved by South Australian Health and Medical Research Institute (SAHMRI) animal ethics committee under application SAM-23-020 and were performed in accordance with ARRIVE guidelines (Percie du Sert et al., 2020). β-ENaC-Tg mice and normal littermates (wild-types) were provided by Charité–Universitätsmedizin Berlin, Berlin, Germany (Mall et al., 2004). Male and female β-ENaC-Tg (n=14) and wild-type (n=13) mice 86-91 days of age were used. All XV imaging and flexiVent lung function tests were performed at the SAHMRI Preclinical Imaging and Research Laboratories (PIRL, South Australia).

### X-ray Velocimetery (XV) imaging

Mice were anaesthetised with a mixture of 75 mg/kg of ketamine (Ceva, Australia) and 1 mg/kg medetomidine (Ilium, Australia), delivered by intraperitoneal (i.p) injection. Once anaesthetised, the mice were prepared by performing a tracheostomy followed by cannulation with a cut-down endotracheal tube (ET; 18 Ga BD Insyte plastic cannula bevel-cut to 15 mm length). The mice were then positioned in an animal holder. All XV imaging was performed in a Permetium preclinical scanner (4DMedical, Australia) (Reyne et al., 2024).

After XV scan acquisition, data were analysed by 4DMedical to determine local specific ventilation throughout the lung. From this, global metrics such as mean specific ventilation (MSV), and ventilation heterogeneity (VH; the interquartile range divided by the mean, which gives a measure of the ventilation variance across the lung) were calculated. Ventilation defect percentage (VDP) describes the percentage of the lung with specific ventilation below a fixed threshold, which defines areas that are thought to be defective. Typically, this threshold is 60% of the mean specific ventilation for that animal/patient (Mathew et al., 2011, Thomen et al., 2015, Thomen et al., 2017). However, changes in mean specific ventilation between animals cause this ‘defective threshold’ to move arbitrarily for each animal, so here we normalise this to use 60% of the mean specific ventilation of the wild-type group to define a fixed threshold and report the normalised ventilation defect percentage (nVDP). To explore regional variations, these parameters were also calculated on subsets of the lung volume. A mask that contained only regions of the lungs at least 3 voxels away from the lung surface in the areas near the rib cage, and from all voxels at the surfaces around the heart cavity was created. This was done by applying a closing filter of kernel size 9×9×9 to a binary lung mask, followed by an erosion filter with size 3×3×3. This mask was used to split the lungs into outer and inner regions, with the outer regions containing 61% of the lung by volume on average. The total lung volume at peak exhalation (including both lung tissue and air) was calculated from the size of the XV volumes produced.

### flexiVent respiratory mechanics

Following the acquisition of XV scans, lung function assessments were conducted using a flexiVent FX small animal ventilator (SCIREQ, Montreal, Canada). The ventilator was equipped with an FX2 mouse module and negative pressure-driven forced expiration (NPFE) extension and operated using flexiWare v8.0 software. Subjects were integrated to the flexiVent via the 18G cannula, and ventilation was performed at rate of 150 breaths/min, an inspiratory ratio of 2:3, tidal volume set at 10 mL/kg, and a positive end expiratory pressure (PEEP) of 3 cmH_2_O. The lung mechanics evaluations were executed utilising SCIREQ’s automated algorithms, which were configured within a mouse mechanics scan script that was repeated three times, as previously described (Devos et al., 2017). Briefly, the script performed a deep inflation manoeuvre, followed by single frequency (SnapShot-150) and broadband (Quick Prime-3) oscillations and a pressure volume loop (PVs-P). A NPFE perturbation was then executed to generate a flow-volume loop and measure parameters such as forced expiratory volume (FEV), forced vital capacity (FVC), and peak expiratory flow (PEF). For each parameter, an average of three measurements was calculated per mouse. Data was excluded if the coefficient of determination – a measure of model fit – was less than 0.9 for each model. After lung analysis mice were humanely killed by i.p overdose of lethabarb (150 mg/kg, Virbac, Australia).

### Tissue analysis

Lungs were inflation-fixed *in situ* using a 10% formalin buffer at 18 cm H_2_O, lungs were then removed and placed into formalin for at least 24 hours and transferred in 70% ethanol until processing. Samples were embedded in paraffin wax and 5 µm slices were mounted onto slides and stained with Alcian Blue/Periodic Acid Schiff for detection of mucus. Images were captured using a Nikon Eclipse Ci-L plus Microscope with a Nikon DS F*i*3 camera and NIS-elements D software (version 5.42.03).

### Statistics

All statistical analyses were performed in R version 4.3.1 (R Core Team, 2022). In keeping with the development of modern bio-statistical analyses that better reveal and describe the clinical relevance of findings – as opposed to providing only binary *p* value decisions about statistical significance – we express our statistical findings in terms of estimated marginal means and 95% confidence intervals returned from linear models fitted to the data. For every flexiVent and XV parameter, a standard linear regression model was fitted using the “lm” function. The model incorporated a fixed effect of genotype and adjustments for age and sex. *Post hoc* pairwise comparisons for the fitted model were carried out using the “emmeans’’ package (Lenth, 2022). Supplementary Table 1 shows the estimated differences in mean between the genotypes and 95% confidence intervals, and the actual *p* value for each parameter.

## Acknowledgements

The authors acknowledge the facilities and scientific and technical assistance of the National Imaging Facility, a National Collaborative Research Infrastructure Strategy (NCRIS) capability, at the Large Animal Research and Imaging Facility, South Australian Health and Medical Research Institute.

## Authors’ Contributions

NR: Conceptualisation, Data Curation, Formal analysis, Investigation, Methodology, Writing - Original Draft.

RS: Data Curation, Formal Analysis, Investigation, Software, Visualization, Writing - Original Draft. PC: Investigation, Data Curation, Formal analysis, Writing - Original draft.

NE, JL, ML, KN: Formal analysis, Writing – Review and editing. JD: Resources, Writing – Review and editing.

MAM, DP: Conceptualisation, Resources, Funding acquisition, Writing – Review and editing.

MD: Conceptualisation, Data Curation, Formal analysis, Funding acquisition, Investigation, Methodology, Supervision, Visualization, Writing - Original Draft.

## Data Availability

The data is hosted in a repository at DOI: 10.25909/25814851

